# H3K79 methylation and H3K36 tri-methylation synergistically regulate gene expression in pluripotent stem cells

**DOI:** 10.1101/2025.05.08.652740

**Authors:** Emmalee W. Cooke, Cheng Zeng, Suza Mohammad Nur, Yunbo Jia, Aileen Huang, Jiwei Chen, Peidong Gao, Fei Xavier Chen, Fulai Jin, Kaixiang Cao

## Abstract

In metazoans, nucleosomes harboring H3K79 methylation (H3K79me) deposited by the histone methyltransferase DOT1L decorate actively transcribed genes. Although DOT1L is implicated in transcription regulation and pathogenesis of human diseases such as leukemia and neurological disorders, the role of H3K79me in these biological processes remains elusive. Here, we reveal a novel functional synergism between H3K79me and H3K36 tri-methylation (H3K36me3), another histone modification enriched at active genes, in regulating gene expression and neural cell fate transition. Simultaneous catalytic inactivation of DOT1L and the H3K36 methyltransferase SETD2 via gene editing leads to the global loss of H3K79me and H3K36me3, hyperactive transcription, and failures in neural differentiation. Interestingly, the loss of H3K79me and H3K36me3 causes increased transcription elongation, gained chromatin accessibility at a group of enhancers, and increased binding of TEAD4 transcription factor and its co-activator YAP1 at these enhancers. Furthermore, YAP-TEAD inhibition partially restores the expression levels of hyperactivated genes upon H3K79me/H3K36me3 loss. Taken together, our study demonstrates a synergistic role of H3K79me and H3K36me3 in regulating transcription and cell fate transition, unveils novel mechanisms underlying such synergism, and provides insight into designing therapies that target diseases driven by misregulation or mutations of DOT1L and/or SETD2.

## Introduction

The nucleosome, the fundamental building block of eukaryotic chromatin, consists of an octamer of four core histones (H2A, H2B, H3, and H4) wrapped by 147 bp of DNA. The tails of core histones are heavily post-translationally modified by histone-modifying enzymes. These histone modifications are believed to instruct essential biological processes such as cell proliferation, differentiation, and organism development via regulating gene transcription; however, recent results suggest that multiple types of histone modifications are dispensable for the regulation of gene expression and cell identity (*1-8*), revealing a missing link between histone modifications and their proposed biological functions.

In mammals, actively transcribed genes are decorated by mono-, di-, and tri-methylation on lysine 79 of histone H3 (H3K79me1/2/3 or H3K79me in general) deposited by the histone methyltransferase DOT1L, which plays a critical role in normal development and human pathologies such as leukemia and neurological disorders (*9-12*). DOT1L interacts with the phosphorylated C-terminal domain (CTD) of RNA polymerase II (RNAPII), suggesting a link between the methyltransferase and transcription elongation (*13*). Indeed, DOT1L cooperates with the super elongation complex to regulate transcription elongation in murine embryonic stem cells (ESCs) (*4*). Although DOT1L is heavily studied in different systems, little is known about the role of H3K79me. Namely, the demethylation pathways of H3K79me and the “reader” proteins that specifically recognize H3K79me on chromatin remain enigmatic. Despite the link between DOT1L and transcription elongation, H3K79me loss has little impact on transcription elongation in ESCs (*4*), suggesting that the catalytic activity is not the major mechanism underlying the transcriptional regulatory role of DOT1L. To date, whether H3K79me plays an instructive role in regulating transcription has remained unclear.

Besides H3K79me, H3K36me3 is another histone mark enriched on the bodies of active genes (*14-17*). In mammals, H3K36me3 is catalyzed by the histone methyltransferase SETD2, the loss of which leads to embryonic lethality and aberrant cortical patterning (*18-20*). Similar to DOT1L, SETD2 binds phosphorylated CTD of RNAPII (*21-23*), depositing H3K36me3 co-transcriptionally on gene bodies. It is proposed that H3K36me3 recruits the *de novo* DNA methyltransferase DNMT3B, which deposits intragenic DNA methylation on active genes and suppresses spurious transcription initiation (*24, 25*). This model suggests that H3K36me3 functions as a repressor to fine-tune transcription fidelity; however, most previous studies utilize knockout or knockdown approaches to target SETD2 and thus are unable to tease apart catalytic-dependent and -independent functions of SETD2.

The colocalization of H3K79me and H3K36me3 on active genes prompted the hypothesis that these two modifications synergize to regulate gene expression. To examine if H3K79me functions together with H3K36me3 to regulate transcription, here we have abolished both H3K79me and H3K36me3 by simultaneously catalytically inactivating DOT1L and SETD2 in ESCs via CRISPR-Cas9 mediated gene editing. Unexpectedly, the loss of H3K79me and H3K36me3 leads to gene hyperactivation in ESCs and failures of neural differentiation although catalytic inactivating individual DOT1L or SETD2 only causes moderate de-regulation in gene expression. Moreover, depleting H3K79me and H3K36me3 results in increased RNAPII elongation and chromatin recruitment of transcription factor TEAD4 and its co-factor YAP1 at enhancers. Furthermore, the gene hyperactivation and increased chromatin accessibility caused by H3K79me/H3K36me3 loss are partially restored by YAP/TEAD inhibition. Our results not only demonstrate a novel synergistic role of H3K79me and H3K36me3 in suppressing gene hyperactivation, but also unveil YAP/TEAD recruitment as a molecular mechanism underlying gene misregulation caused by H3K79me/H3K36me3 loss, suggesting alternative strategies to target diseases driven by DOT1L and/or SETD2 loss-of-function.

## Results

### Genome-wide colocalization of H3K79me and H3K36me3

We previously demonstrated that H3K79me is largely dispensable for ESC self-renewal and neural differentiation (*4*); however, catalytic inactivation of DOT1L leads to embryonic lethality in mice (*26*), suggesting that H3K79me is essential for mammalian development. To date, the role of H3K79me in regulating transcription and cell fate transition remains elusive. We surmise that H3K79me may function through the crosstalk with other epigenetic pathways. In metazoans, active genes are decorated with H3K36me3 in addition to H3K79me (*14-17*). To compare the genome-wide occupancy of H3K79me and H3K36me3, we performed chromatin immunoprecipitation with reference exogenous genome (ChIP-Rx) analysis in wildtype (WT) ESCs with antibodies against H3K79me1, H3K79me2, H3K79me3, and H3K36me3. Our analyses indicated that H3K79me1/2/3 (hereafter represented by H3K79me) and H3K36me3 were enriched on gene bodies in ESCs (Fig. S1A and Fig. 1A-C). Consistent with previous reports (*16, 17, 27*), such enrichment was positively correlated with the levels of gene expression (Fig. S1B and Fig. 1D-F). As expected, peaks of H3K79me1, H3K79me2, and H3K79me3 had a major overlap (Fig. 1G). Interestingly, 64% of H3K79me1 peaks, 61% of H3K79me2 peaks, and 64% of H3K79me3 peaks colocalized with H3K36me3 peaks (Fig. S1C-E), although H3K79me was more enriched at the 5’ of genes and H3K36me3 was more enriched at the 3’ of genes. Moreover, ∼63% of H3K36me3 peaks overlapped with merged H3K79me1/2/3 peaks, and genes under the 28,934 H3K79me/H3K36me3 overlapping peaks had significantly higher expression levels than genes outside these peaks (Fig. 1H-I, Fig. S1F-H). These results suggest a potential functional synergism between H3K79me and H3K36me3 in regulating active genes.

**Figure 1.**
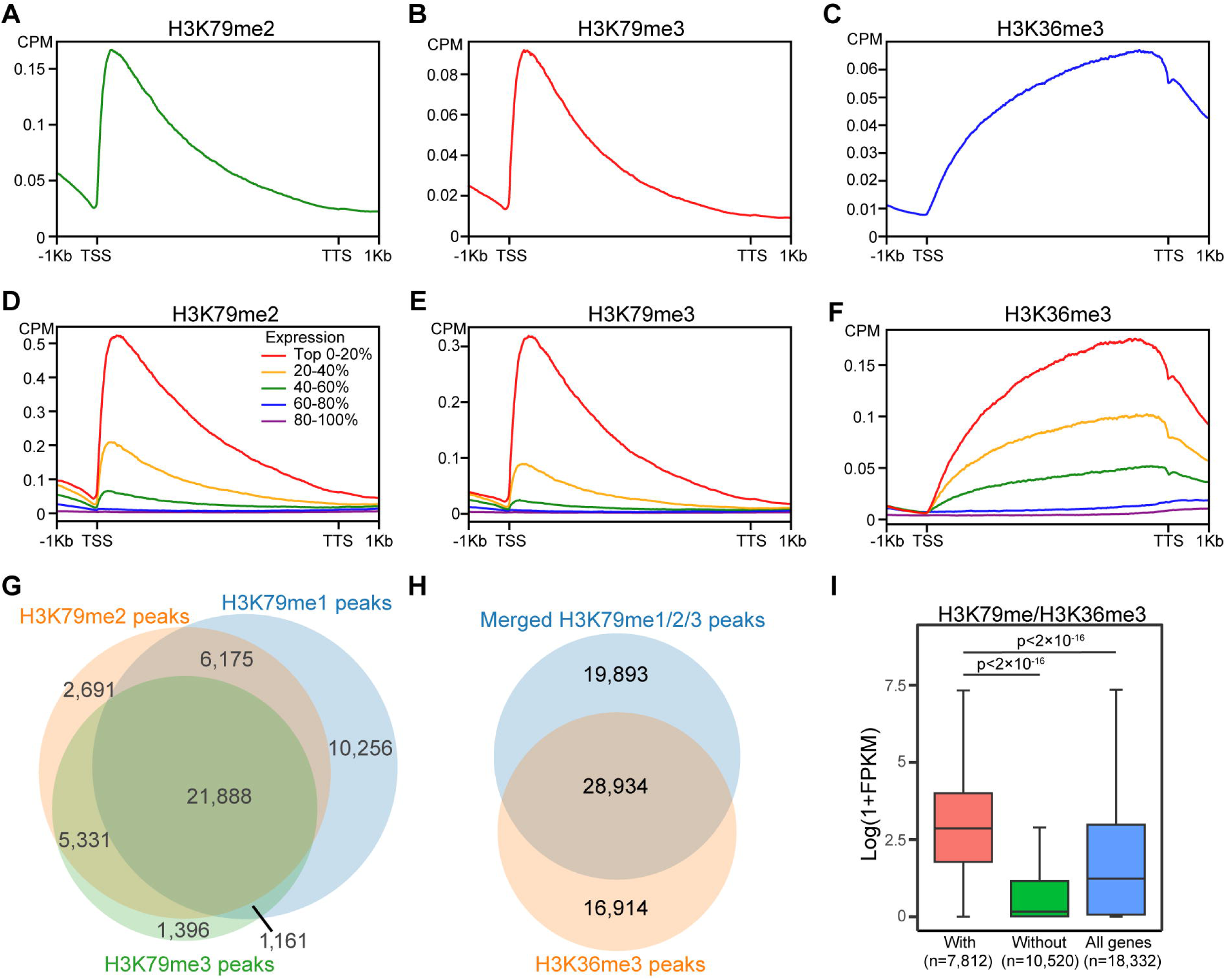
H3K79me and H3K36me3 are enriched at active genes. (A-C) Metaplots of average ChIP-Rx signals of H3K79me2 (A), H3K79me3 (B), and H3K36me3 (C) on all protein-coding genes (n=18,332) in ESCs. CPM: spike-in normalized counts per million reads. TSS: transcription start site; TTS: Transcription termination site. (D-F) Metaplots showing the signals of H3K79me2 (D), H3K79me3 (E), and H3K36me3 (F) on all genes categorized into five groups based on their expression levels. (G) Venn diagram showing the colocalization between enriched regions of H3K79me1, H3K79me2, and H3K79me3. (H) Venn diagram showing the colocalization of H3K79me1/2/3 merged peaks and H3K36me3 peaks. (I) Box plot analysis of the expression levels of genes with and without H3K79me/H3K36me3 peaks. The number of genes in each category is labeled.

### Simultaneous catalytic inactivation of DOT1L and SETD2 leads to the global loss of H3K79me and H3K36me3

To understand whether H3K79me and H3K36me3 co-regulate gene expression, we deleted the majority of the <u>S</u>u(var)3-9, <u>E</u>nhancer-of-zeste, and <u>T</u>rithorax (SET) domain of SETD2, which catalyzes H3K36me3 (*28, 29*), in both WT and DOT1L catalytic-inactive (CI) ESCs(*4*) using CRISPR-Cas9 guided genetic engineering to generate respective SETD2 CI and DOT1L/SETD2 double CI (DCI) cells (Fig. 2A). RNA-seq analysis indicated that the targeted exons 6-8 of the *Setd2* gene were deleted in SETD2 CI and DCI cells (highlighted in Fig. S2A). The in-frame deletion did not impact the level of SETD2 protein in nucleoplasm or on chromatin (Fig. S2B-C), suggesting that the level of SETD2 and its recruitment to chromatin is not impaired by the loss of its SET domain or catalytic activity. Alkaline phosphatase staining and RNA-seq analysis indicated that the morphology and expression levels of pluripotency genes of SETD2 CI and DCI cells were comparable to those of WT and DOT1L CI cells (Fig. 2B and Fig. S2D), suggesting that the self-renewing capability of ESCs is not impaired by the catalytic inactivation of DOT1L or/and SETD2. To determine if deleting the SET domain of SETD2 impairs H3K36me3 levels in ESCs, we examined the levels of H3K36me3 with histones extracted from WT, DOT1L CI, SETD2 CI, and DCI cells by Western blotting. DOT1L CI cells harbored comparable levels of H3K36me3 to WT cells while H3K36me3 was undetectable in SETD2 CI and DCI cells (Fig. 2C). We further showed that DOT1L and/or SETD2 inactivation had little impact on global H3K36me2 level (Fig. S2F), indicating the specificity of SETD2 in depositing H3K36me3 in ESCs. As expected, Western blotting analysis indicated a loss of H3K79me2 and H3K79me1 in both DOT1L CI cells and DCI cells (Fig. 2D and Fig. S2E). We further performed H3K36me3, H3K79me1, H3K79me2, and H3K79me3 ChIP-Rx in WT, DOT1L CI, SETD2 CI, and DCI cells. As expected, H3K79me and H3K36me3 occupancies correlated with RNAPII levels on protein-coding genes (Fig. 2E-I). Our analyses indicated that H3K36me3 and H3K79me were abolished at all protein-coding genes in SETD2 CI cells and DOT1L CI cells, respectively (Fig. 2F-I and Fig. S2G). Furthermore, neither H3K36me3 nor H3K79me were detectable at protein-coding genes in DCI cells. Interestingly, we observed a slight decrease in H3K36me3 levels toward the 3’ of genes in DOT1L CI compared with WT cells (Fig. S2H), while the levels of H3K79me1/2/3 all had a moderate increase upon H3K36me3 loss (Fig. S2I-K), suggesting a crosstalk between H3K36me3 and H3K79me. Taken together, our data indicate that the simultaneous catalytic inactivation of DOT1L and SETD2 leads to a global loss of H3K79me and H3K36me3 and that ESC self-renewal is not impaired by the loss of H3K79me and H3K36me3.

**Figure 2.**
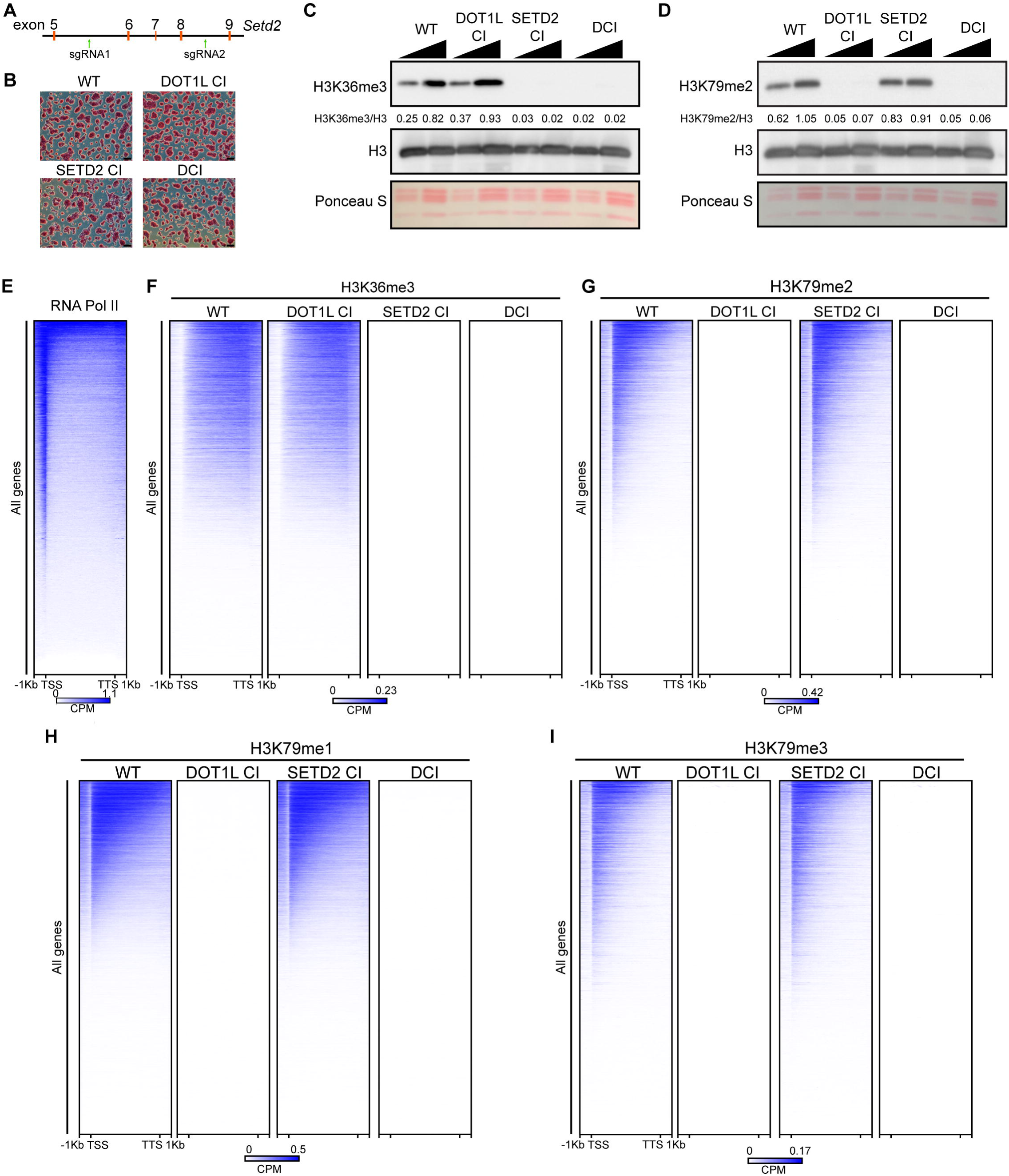
Simultaneous catalytic inactivation of DOT1L and SETD2 depletes H3K79me and H3K36me3 globally. (A) Schematic representation of the localizations of two sgRNAs for deleting the SET domain of the *Setd2* gene. (B) Alkaline phosphatase staining of WT, DOT1L CI, SETD2 CI, and DCI ESCs. Scale bar: 100μm. (C-D) Western blots indicating the level of H3K36me3 (C) and H3K79me2 (D) in WT, DOT1L CI, SETD2 CI, and DCI ESCs. 1 and 2 μg extracted histones were loaded for each sample. The levels of histone modifications in each lane were quantified by normalizing with H3 signals of the corresponding lane. Normalized ratios were provided under the modification blot. (E) Heatmap showing RNAPII levels on all protein-coding genes (n=18,332) in ESCs. Genes are sorted by descending occupancy of RNAPII at TSSs in WT ESCs. (F-I) Heatmaps of H3K36me3 (F), H3K79me2 (G), H3K79me1 (H), and H3K79me3 (I) levels at all protein-coding genes (n=18,332) in WT, DOT1L CI, SETD2 CI, and DCI ESCs. Genes are sorted the same way as in (E).

### H3K79me and H3K36me3 synergistically regulate gene expression

To examine the role of H3K79me and H3K36me3 in regulating gene expression, we performed RNA-seq using WT, DOT1L CI, SETD2 CI, and DCI ESCs. Our analysis indicated that the catalytic inactivation of DOT1L or SETD2 caused moderate changes in gene expression (320 significantly differentially expressed genes in DOT1L CI and 190 in SETD2 CI cells) with more upregulated than downregulated genes (Fig. 3A-B). Interestingly, the simultaneous loss of H3K79me and H3K36me3 led to the significant misregulation of 1,993 genes in ESCs (Fig. 3C), indicating a synergism between H3K79me and H3K36me3 in regulating gene expression. To our surprise, H3K79me/H3K36me3 loss led to the significant upregulation of 1,418 genes, accounting for more than 70% of misregulated genes in DCI cells (Fig. 3C). Overlapping and gene set enrichment analysis (GSEA) showed that misregulated genes in DOT1L CI and SETD2 CI cells were significantly deregulated in DCI cells (Fig. S3A-C). On the other hand, 80% of upregulated genes in DCI cells (1,135 of 1,418) including highly expressed genes such as *Gsk3b* and *Abcc5* were not significantly upregulated in DOT1L CI or SETD2 CI cells (Fig. S3A and Fig. 3D), further indicating the synergism between H3K79me and H3K36me3 in dampening gene expression.

**Figure 3.**
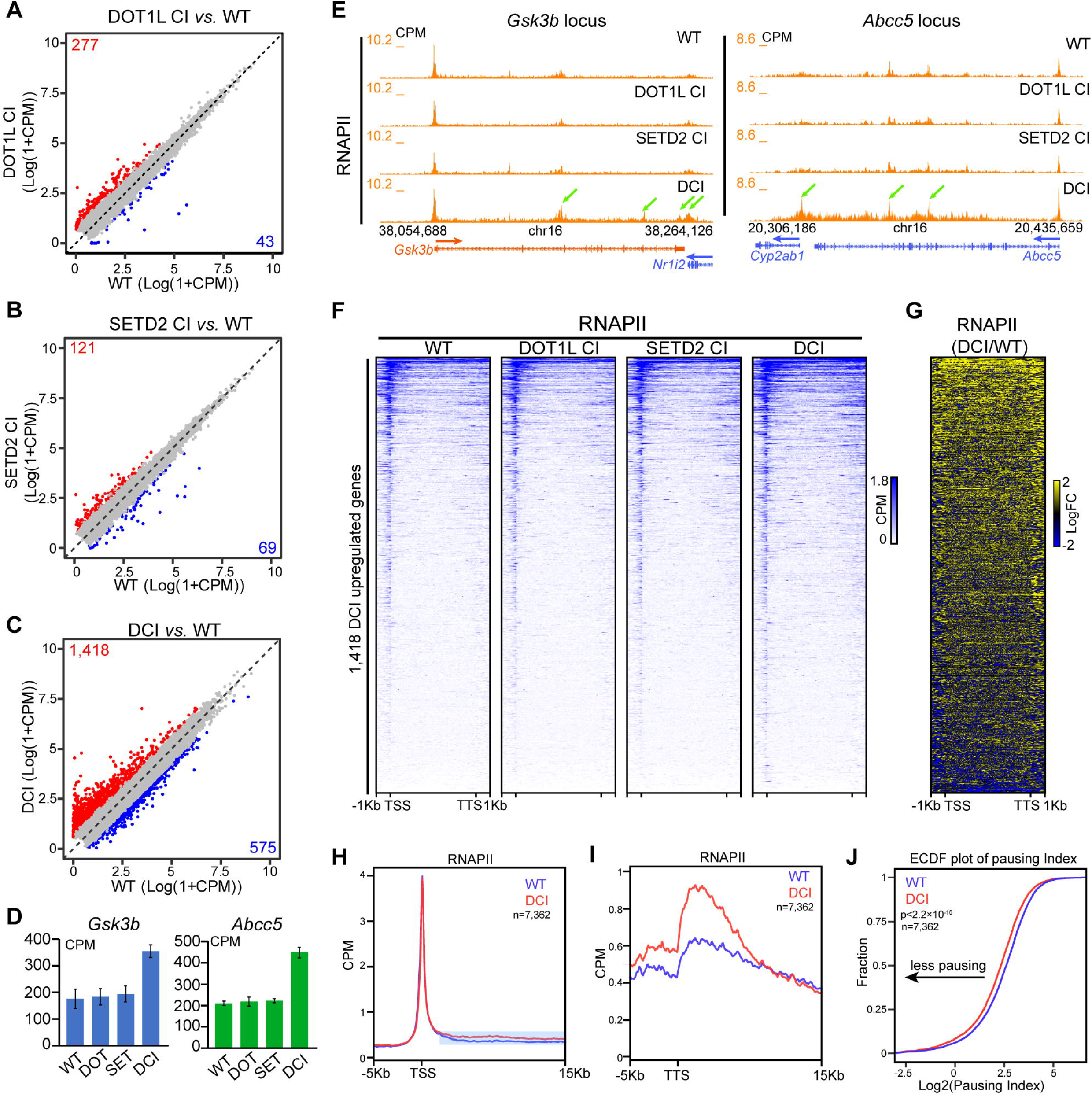
The loss of H3K79me and H3K36me3 leads to transcription hyperactivation. (A-C) Correlation plots of RNA-seq data between DOT1L CI (A), SETD2 CI (B), or DCI (C) cells and WT cells. Statistical significance was determined by a two-sided Wald test and p-values were corrected for multiple testing using the Benjamini-Hochberg method. Significantly upregulated genes (log2 fold change >1, adjusted p<0.01) are highlighted in red while downregulated (log2 fold change <-1, adjusted p<0.01) genes are highlighted in blue. The number of up- and downregulated genes are listed on the plots. (D) RNA-seq signals of *Gsk3b* (left) and *Abcc5* (right) genes in WT, DOT1L CI, SETD2 CI, and DCI cells. (E) Genome browser view of RNAPII ChIP-Rx data at the *Gsk3b* (left) and *Abcc5* (right) loci in WT, DOT1L CI, SETD2 CI, and DCI ESCs. (F) Heatmaps showing RNAPII occupancy at 1,418 DCI upregulated genes in WT, DOT1L CI, SETD2 CI, and DCI ESCs. Genes are sorted by descending occupancy of RNAPII at TSSs. (G) Heatmap showing the log2 fold change of RNAPII occupancy at 1,418 DCI upregulated genes comparing DCI and WT cells. Genes are sorted the same way as in (F). (H-I) Metaplot analysis of RNAPII levels at -5Kb and +15Kb around TSS (H) and TTS (I) in WT and DCI ESCs on 7,362 RNAPII positive genes previously defined (*4*). Regions downstream of TSSs with increased RNAPII upon the H3K79me/H3K36me3 loss are highlighted. (J) Empirical cumulative density function (ECDF) plots of RNAPII pausing indexes in WT and DCI ESCs at the 7,362 RNAPII positive genes. Pausing indexes were calculated through dividing the RNAPII level at promoters by that on gene bodies. Promoters are defined as the -200 to +400bp regions relative to TSS. Gene bodies are defined as the +400bp from TSS to TTS.

To determine whether the upregulated genes in DCI cells are transcriptionally activated, we performed RNAPII ChIP-Rx in WT, DOT1L CI, SETD2 CI, and DCI cells. Our data indicated that the levels of RNAPII were increased at the 1,418 DCI upregulated genes in DCI cells compared with WT cells; however, RNAPII levels at these genes in DOT1L CI and SETD2 CI cells were comparable to that in WT cells (Fig. 3E-G). We noted that the H3K79me/H3K36me3 loss led to the gain of RNAPII levels on the bodies of DCI upregulated genes (Fig. 3E-G, Fig. S3D), suggesting that H3K79me and H3K36me3 dampen transcription elongation in a synergistic manner. On the other hand, serine 5 phosphorylated RNAPII (Ser5P) levels on DCI upregulated genes *Gsk3b* and *Abcc5* were not increased (Fig. S3E), suggesting that the gained RNAPII at DCI upregulated genes is unlikely due to increased spurious transcription initiation, which has been shown to be a consequence of SETD2 depletion (*18, 25*). To further examine if the simultaneous loss of H3K79me and H3K36me3 has an impact on transcription elongation, we compared RNAPII levels in WT and DCI cells at promoters and gene bodies of 7,362 RNAPII positive genes (*4*). H3K79me/H3K36me3 loss led to increased RNAPII levels downstream of transcription start sites (TSSs) while the RNAPII levels at TSSs in WT and DCI cells were comparable (Fig. 3H). Moreover, RNAPII levels at transcription termination sites (TTSs) were elevated in DCI compared with WT cells (Fig. 3I). We further calculated the pausing index at RNAPII positive genes in WT and DCI cells based on the ratio of the level of RNAPII at promoters (-200bp to +400bp relative to TSS) and that on gene bodies (from 400 bp downstream of TSS to TTS). Empirical cumulative density function (ECDF) analysis indicated that pausing indexes at RNAPII positive genes were decreased in DCI compared with WT cells (Fig. 3J), suggesting that the loss of H3K79me and H3K36me3 leads to increased transcription elongation. Furthermore, ChIP-Rx analysis indicated that serine 2 phosphorylated RNAPII (Ser2P) was increased at TTSs of the 1,418 DCI upregulated genes and 7,362 RNAPII positive genes (Fig. S3F-G), suggesting the elevated transcription elongation caused by H3K79me/H3K36me3 loss. Taken together, these results indicate that H3K79me and H3K36me3 synergistically suppress gene hyperactivation and dampen transcription elongation.

### H3K79me and H3K36me3 synergistically facilitate neural lineage specification

We previously demonstrated that DOT1L is required for neural lineage specification; however, H3K79me loss does not impair glial or neuronal differentiation (*4*), leaving the function of H3K79me in cell fate transition largely unknown. On the other hand, SETD2 deletion in neural precursor cells causes abnormal area patterning of the cerebral complex in mice (*20*), pointing to a potential role of H3K36me3 in neural development. To determine if H3K79me and H3K36me3 synergize to regulate neural differentiation, we utilized a well-established neural differentiation protocol (*4*) to differentiate WT, DOT1L CI, SETD2 CI, and DCI ESCs toward astrocytes and neurons. We noticed that DCI ESCs generated smaller embryoid bodies at day 6 of the neural differentiation scheme compared with WT, DOT1L CI, or SETD2 CI cells (Fig. S4A-B), suggesting a synergism between H3K79me and H3K36me3 in regulating cell fate transition. Although H3K79me loss had little impact on glial and neuronal specification (Fig. S4C and (*4*)), H3K36me3 depletion impaired neural differentiation as fewer neurons and astrocytes were found after 29 days of differentiation (Fig. S4C), consistent with the role of SETD2 in neural development. Interestingly, the loss of H3K79me/H3K36me3 in ESCs completely blocked neuronal and glial specification (Fig. S4C), suggesting that H3K79me and H3K36me3 regulate neural lineage commitment in a synergistic manner.

To determine the mechanisms underlying the impact of H3K79me/H3K36me3 loss on neural differentiation, we performed RNA-seq with the WT and DCI cultures at day 29 of neural differentiation. 3,419 differentially expressed genes were identified in DCI cultures compared with their WT counterpart (Fig. S4D), indicating a drastic impact of H3K79me/H3K36me3 loss on the transcriptome during neural differentiation. As expected, gene ontology (GO) analysis indicated that genes related to neural development are enriched in significantly downregulated genes in DCI compared with WT cultures (Fig. S4E). Moreover, marker genes of neural cells were significantly downregulated in DCI cultures (Fig. S4G), corroborating results that neuronal and glial specification are blocked in DCI cells. To our surprise, adipogenesis-related genes were significantly enriched in genes upregulated by H3K79me/H3K36me3 loss (Fig. S4F). Consistently, marker genes of adipogenesis were significantly upregulated in DCI cells after neural differentiation (Fig. S4G), suggesting that the H3K79me and H3K36me3 could safeguard neural lineage commitment by suppressing the hyperactivation of adipogenesis-related genes. Taken together, our data demonstrate that H3K79me and H3K36me3 synergistically facilitate neuronal and glial specification and the suppression of adipogenesis-related genes during neural differentiation.

### Increased recruitment of YAP-TEAD to enhancers decompacted by the H3K79me/H3K36me3 loss

To further dissect the mechanism underlying gene hyperactivation caused by the simultaneous loss of H3K79me and H3K36me3, we performed the assay for transposase-accessible chromatin using sequencing (ATAC-seq) in WT and DCI ESCs and identified 2,089 significantly increased ATAC-seq peaks in DCI ESCs (Fig. S5A-B). Besides enriched at active gene bodies, both H3K79me and H3K36me3 are found at enhancers in various models (*30-34*), suggesting that these modifications could contribute to enhancer regulation. Indeed, more than 80% of the increased ATAC-seq peaks in DCI cells were located more than 5Kb away from TSS (Fig. S5C). Moreover, levels of H3K36me3 and H3K79me were significantly reduced at increased ATAC-seq peaks in DCI cells upon DOT1L/SETD2 catalytic inactivation (Fig. S5D-G), suggesting that the loss of H3K79me/H3K36me3 leads to chromatin decompaction at these enhancers. We then examined the levels of enhancer marks H3K27ac and H3K4me1 at increased ATAC-seq peaks in WT and DCI cells by ChIP-Rx analysis. The levels of H3K27ac and H3K4me1 were increased at regions that gained accessibility upon H3K79me/H3K36me3 loss (Fig. 4A-C). Moreover, genes near these ATAC-seq peaks were significantly upregulated in DCI compared with WT cells (Fig. 4D). Furthermore, H3K27ac and H3K4me1 were increased at multiple enhancers of H3K79me/H3K36me3 target genes *Gsk3b* and *Abcc5* (Fig. 4E-F). These results suggest that the loss of H3K79me and H3K36me3 leads to the decondensation and hyperactivation of a group of enhancers, which correlates with gene upregulation observed in DCI cells.

**Figure 4.**
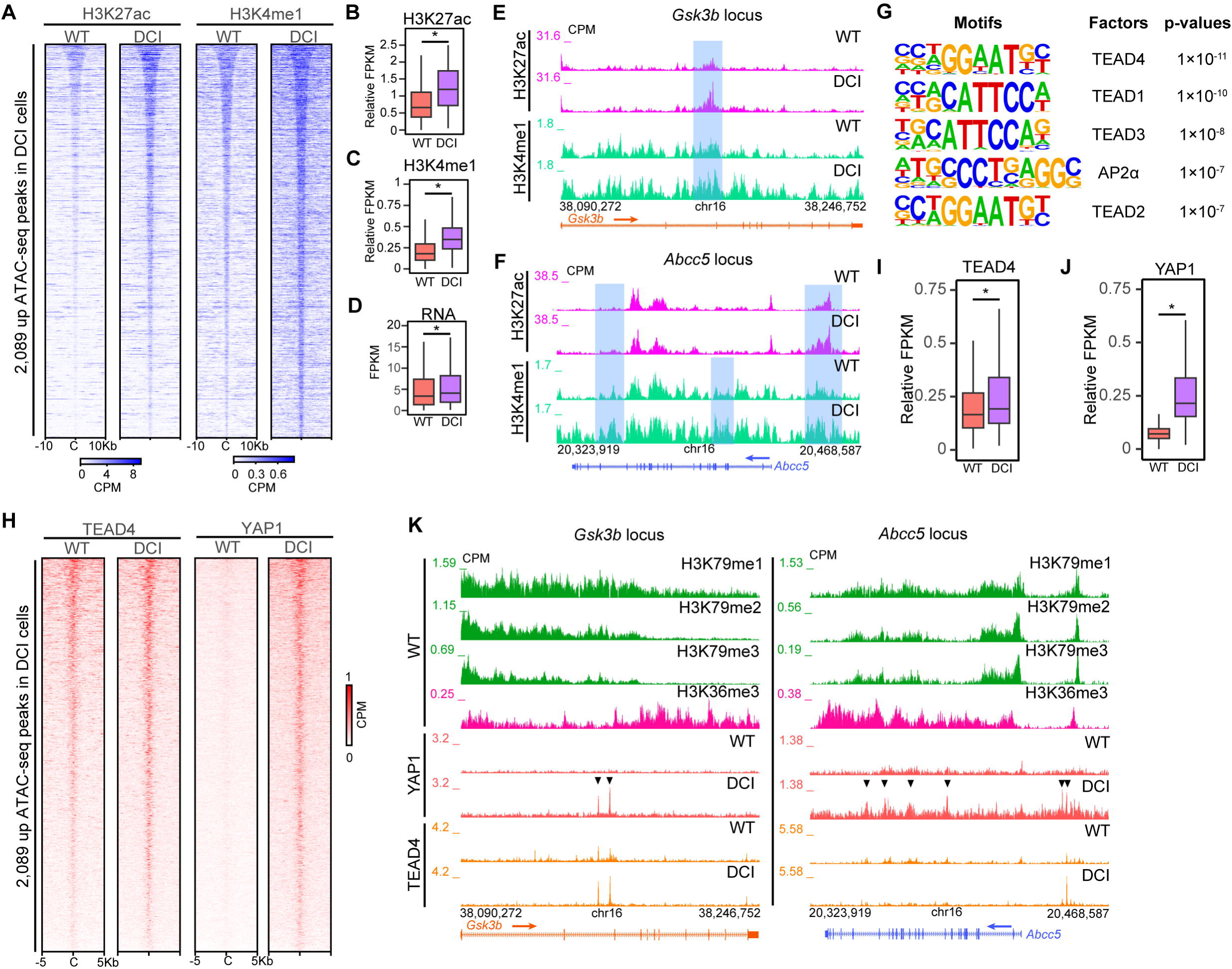
The loss of H3K79me and H3K36me3 leads to an increased recruitment of YAP-TEAD to enhancers. (A) Heatmaps showing the levels of H3K27ac (left) and H3K4me1 (right) at DCI-upregulated ATAC-seq peaks in WT and DCI cells. Data are sorted by descending ATAC-seq signals in WT cells. (B-C) Box plot analysis indicating the levels of H3K27ac (B) and H3K4me1 (C) at DCI-upregulated ATAC-seq peaks in WT and DCI cells. *: p<2×10^-16^. (D) Box plot analysis indicating the expression levels of nearest genes to DCI-upregulated ATAC-seq peaks in WT and DCI cells. *: p<2×10^-16^. (E-F) Genome browser view of H3K27ac and H3K4me1 ChIP-Rx data in WT and DCI cells at *Gsk3b* (E) and *Abcc5* (F) loci. Regions that gain H3K27ac and/or H3K4me1 in DCI cells are highlighted in blue. (G) Top transcription factor motifs enriched under upregulated ATAC-seq peaks in DCI ESCs. (H) Heatmaps showing the levels of TEAD4 and YAP1 at DCI-upregulated ATAC-seq peaks in WT and DCI cells. Data are sorted by descending ATAC-seq signals in WT cells. (I-J) Box plot analysis indicating the levels of TEAD4 (I) and YAP1 (J) at DCI-upregulated ATAC-seq peaks in WT and DCI cells. *: p<2×10^-16^. (K) Genome browser view of YAP1 and TEAD4 ChIP-Rx data in WT and DCI cells at *Gsk3b* (left) and *Abcc5* (right) loci. H3K79me1, H3K79me2, H3K79me3, and H3K36me3 ChIP-Rx data in WT cells at these two loci are also shown. Arrows indicate regions that gain YAP1 in DCI cells.

To unveil why specific enhancers are hyperactivated in DCI cells, we performed motif analysis at increased ATAC-seq peaks upon the loss of H3K79me and H3K36me3. Our analysis indicated that TEAD transcription factors were significantly enriched at these peaks (Fig. 4G), suggesting that the occupancy of TEADs is increased at these regions in DCI cells. Indeed, ChIP-Rx analysis indicated that the levels of TEAD4 and its co-factor YAP1 were significantly increased at DCI-upregulated ATAC-seq peaks upon H3K79me/H3K36me3 loss (Fig. 4H-J). For instance, enhancers under H3K79me and H3K36me3 enriched regions at *Gsk3b* and *Abcc5* harbored increased YAP1 and TEAD4 occupancy in DCI compared with WT ESCs (Fig. 4K). YAP1 and TEADs are critical components of the Hippo signaling pathway (*35, 36*). Phosphorylated YAP1 is sequestered in the cytoplasm while unphosphorylated YAP1 enters the nucleus, binds TEADs, and is recruited to chromatin to activate the transcription of TEAD target genes (*37-41*). To determine if the catalytic activity of DOT1L and SETD2 plays a role in regulating the Hippo signaling pathway, we examined the levels of YAP1 and phosphorylated YAP1 in WT and DCI cells. Western blotting analysis indicated that catalytic inactivation of DOT1L and SETD2 has little impact on the levels of YAP1 or phosphorylated YAP1 (Fig. S6A-B). Furthermore, cytoplasmic and nuclear levels of YAP1 were comparable in WT and DCI cells (Fig. S6C-D), indicating that the phosphorylation cascade in the Hippo pathway is unlikely to be impaired by the catalytic inactivation of DOT1L and SETD2. Overall, these results suggest that the loss of H3K79me/H3K36me3 could facilitate the recruitment of YAP-TEAD to its target enhancers, which may lead to gene hyperactivation in DCI cells.

### YAP-TEAD inhibition rescues gene hyperactivation caused by the H3K79me/H3K36me3 loss

To unveil if the increased binding of YAP-TEAD on chromatin contributes to gene hyperactivation upon the loss of H3K79me/H3K36me3, we treated DCI cells with verteporfin (VP), which disrupts the interaction between YAP and TEAD (*42, 43*). As expected, 8-hour VP treatment led to the decrease of YAP1 levels in DCI ESCs (Fig. 5A). Moreover, YAP-TEAD inhibition in DCI cells caused the time-dependent downregulation of *Gsk3b* and *Abcc5* (Fig. 5B-C), two genes hyperactivated by the loss of H3K79me/H3K36me3. Importantly, GSEA and hierarchical clustering analysis indicated that upregulated genes in DCI cells were significantly downregulated by VP treatment (Fig. 5D-E, Fig. S6E), strongly suggesting that the increased chromatin accessibility of YAP-TEAD at its target enhancers contributes to gene hyperactivation in DCI cells. Although the impact of VP treatment on downregulated genes in DCI cells was not significant at the 8-hour timepoint, these genes were significantly upregulated upon VP treatment for 24 hours (Fig. S6F-G), suggesting that the effect of VP on DCI-downregulated genes could be secondary to that on DCI-upregulated genes. To examine whether YAP-TEAD inhibition could restore chromatin accessibility upon the H3K79me/H3K36me3 loss, we further performed ATAC-seq using VP- and DMSO-treated DCI cells, respectively. YAP-TEAD inhibition led to the decrease of ATAC-seq signals at DCI upregulated genes *Gsk3b* and *Abcc5* (Fig. 5F-G), suggesting that the recruitment of YAP-TEAD facilitates chromatin decompaction at genes hyperactivated by the loss of H3K79me/H3K36me3. Indeed, VP treatment led to a decrease in ATAC-seq signals at peaks that gain accessibility upon H3K79me/H3K36me3 depletion (Fig. 5H-I). Taken together, these results suggest that the increased recruitment of YAP-TEAD to enhancers is partially responsible for the gene hyperactivation and the gain in chromatin accessibility caused by the H3K79me/H3K36me3 loss.

**Figure 5.**
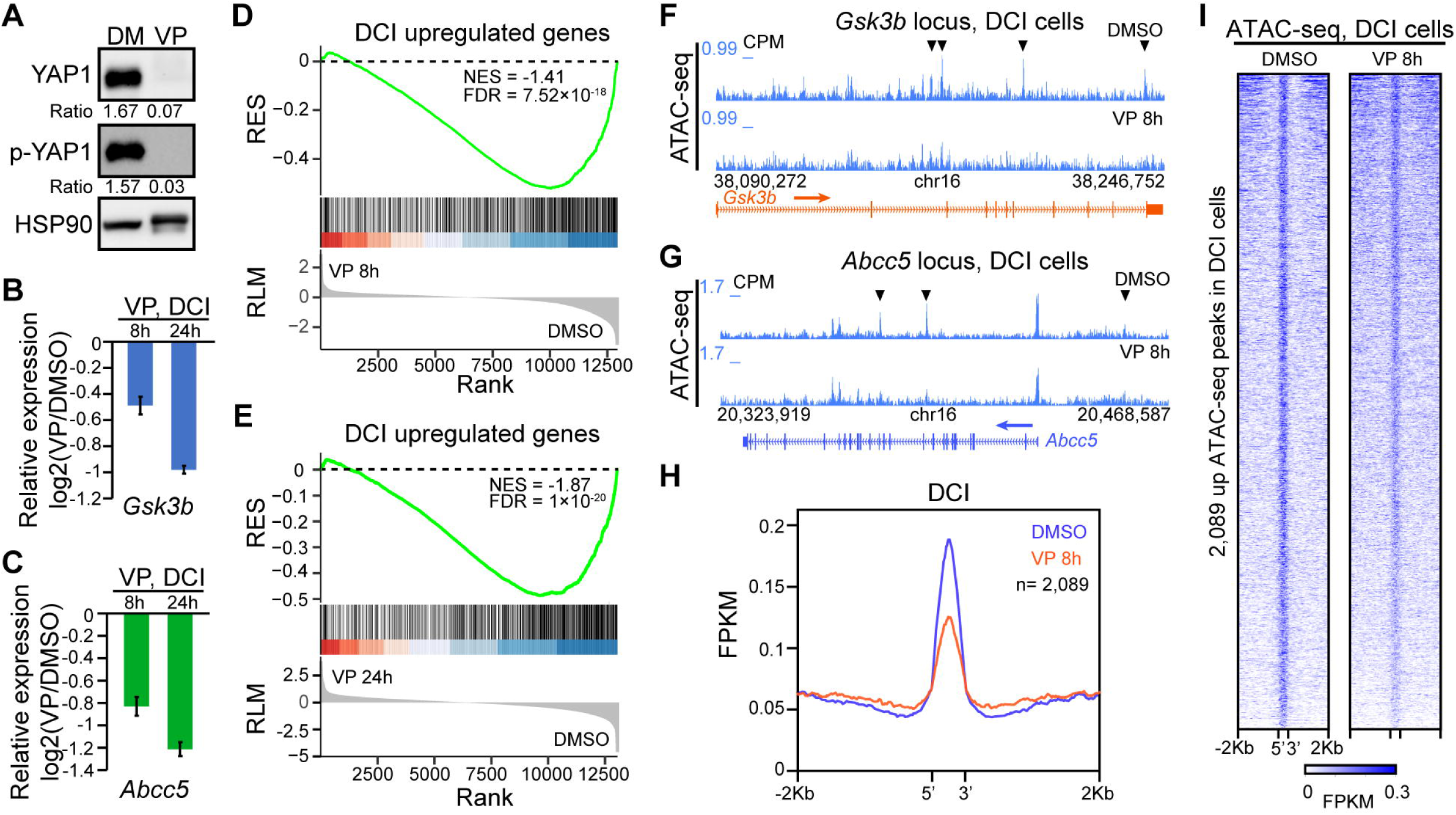
YAP-TEAD inhibition partially rescues the gene misregulation caused by the H3K79me/H3K36me3 loss. (A) Western blotting of YAP1 and phosphorylated-YAP1 (p-YAP1) in DCI cells treated with DMSO or 2 µM verteporfin (VP) for 8 hours. (B-C) Relative expression levels of *Gsk3b* (B) and *Abcc5* (C) in DCI cells treated with 2 µM verteporfin for 8 and 24 hours, respectively. (D-E) GSEA analysis of genes upregulated in DCI cells (red genes in Fig. 3C) comparing 8-hour (D) and 24-hour (E) VP and DMSO treated DCI cells. RES: running enrichment score; RLM: ranked list metric; NES: normalized enrichment score; FDR: false discovery rate. (F-G) Genome browser view of ATAC-seq data in DCI cells treated with DMSO or 2 µM VP for 8 hours at *Gsk3b* (F) and *Abcc5* (G) loci. Arrows indicate regions that gain accessibility upon VP treatment. (H) Metaplot analysis of ATAC-seq levels at DCI-upregulated ATAC-seq peaks in DCI cells treated with DMSO or 2 µM VP for 8 hours. (I) Heatmaps showing ATAC-seq levels at DCI-upregulated ATAC-seq peaks in DCI cells treated with DMSO or 2 µM VP for 8 hours. Data are sorted by descending ATAC-seq signals in DMSO treated DCI cells.

## Discussion

Although the role of DOT1L in regulating gene expression, cell fate, and human diseases such as leukemia has been extensively studied, H3K79me depletion via DOT1L catalytic inactivation in ESCs has little impact on transcription elongation and neural lineage commitment (*4*), leaving the biological functions of H3K79me in gene regulation and cellular differentiation elusive. Here we demonstrated a novel synergistic role of H3K79me and H3K36me3 in dampening hyperactive transcription. The simultaneous loss of H3K79me and H3K36me3 in ESCs leads to increased transcription elongation and the abolishment of neural differentiation. We further showed that the H3K79me/H3K36me3 loss causes increased recruitment of Hippo signaling components YAP1 and TEAD4 to accessible chromatin, which is partially responsible for transcription hyperactivation caused by the loss of H3K79me and H3K36me3.

H3K79me and H3K36me3 are both enriched on the bodies of actively transcribed genes (*14-17*) (Fig. 1), and both DOT1L and SETD2 interact with elongating RNAPII (*13, 21, 22*), suggesting a possible crosstalk and/or synergism between the two modifications. Indeed, our results indicate a crosstalk between H3K79me and H3K36me3: H3K36me3 depletion causes a mild increase in all three types of H3K79me at the 5’ of genes while H3K79me loss leads to a moderate decrease in H3K36me3 levels on the 3’ of genes (Fig. 2 and Fig. S2). Such result is consistent with previous reported H3K79me2 gain on gene bodies upon *Setd2* depletion in leukemia cells (*44*). It is possible that H3K36me3 regulates the spreading of H3K79me on gene bodies and/or the activity of DOT1L on chromatin. The activity of DOT1L on nucleosomes is also regulated by H2BK120 ubiquitination (H2B-Ub), H4K16ac, and the tail of H4 (*45-53*). We surmise that H3K36me3 may indirectly regulate the activity of DOT1L by impacting H2B-Ub, H4K16ac, and/or nucleosome conformation. On the other hand, H3K79me depletion leads to a moderate decrease in H3K36me3 levels at the 3’ of genes. Such decrease may contribute to a higher number of misregulated genes in DOT1L CI cells than that of SETD2 CI cells. Future studies on identifying “readers” and “erasers” of H3K79me would be important to unveil the mechanism underlying the cross talk between H3K79me and H3K36me3.

To our surprise, the loss of H3K79me and H3K36me3 by simultaneously inactivating DOT1L and SETD2 leads to gene hyperactivation (Fig. 3). Increased gene expression in DCI cells is correlated with gained RNAPII levels on gene bodies, suggesting an increased transcription elongation at genes upregulated by H3K79me/H3K36me3 loss. Indeed, we observed increased total RNAPII and Serine 2-phosphorylated RNAPII levels at TTSs and decreased pausing indexes at RNAPII positive genes in DCI compared with WT cells. The increase of RNAPII and elongating RNAPII at TTSs suggests increased transcription beyond the major transcription elongation checkpoint located near the Poly(A) sites (*54*), which may have an impact on transcription termination and 3’ transcript processing. The decreased pausing index in DCI cells is unlikely a result of increased spurious transcription initiation in gene bodies caused by the loss of H3K36me3 as we did not find accumulating serine 5-phosphorylated RNAPII at DCI upregulated genes in DCI compared to WT cells. Interestingly, depletion of ZMYND11, a known “reader” of H3K36me3 (*55, 56*), has been shown to increase transcription elongation. It is possible that H3K79me and H3K36me3 synergistically promote the recruitment of “reader” proteins such as ZMYND11 to suppress transcription elongation. It is noteworthy that the synergism of H3K79me and H3K36me3 is not limited in dampening gene hyperactivation as DCI ESCs have an increased number of significantly downregulated genes compared with DOT1LCI or SETD2 CI cells. To date, menin, a component of the mammalian COMPASS (<u>Com</u>plex of <u>P</u>roteins <u>As</u>sociated with <u>S</u>et1) complex that harbors H3K4 methyltransferase activity (*57*), is the only identified “reader” protein of H3K79me (*58*). It is important to investigate whether H3K79me and H3K36me3 co-activate gene expression via menin, COMPASS, and other potential “readers” of H3K79me and H3K36me3 in future studies to decipher mechanisms underlying the functional synergism of H3K79me and H3K36me3 in gene regulation.

Our results further point to a synergistic role of H3K79me and H3K36me3 in regulating neural differentiation (Fig. S4). The simultaneous loss of H3K79me and H3K36me3 completely blocks the derivation of neuronal and glial cells. Interestingly, adipogenesis-related genes were hyperactivated in DCI cells during neural differentiation. Although adipocytes are thought to derive from mesoderm, they arise from diverse lineages that are heterogeneously distributed (*59*). Besides mesenchymal progenitors, it has been shown that cells from neuroectoderm such as neural crest cells are also capable of differentiating into adipocytes both in culture and *in vivo* (*60, 61*). It is possible that H3K79me and H3K36me3 synergistically suppress the expression of adipogenesis-related genes and adipogenesis from neural crest cells during neural differentiation to safeguard neuronal and glial specification. We surmise that the skewed transcription elongation upon the loss of H3K79me and H3K36me3 contributes to the potential cell fate switch between neural lineage commitment and adipogenesis. Understanding how H3K79me and H3K36me3 synergistically regulate transcription elongation, neural differentiation, and adipogenesis in future studies is important to understand the mechanism underlying disease pathogenesis driven by misregulation and mutations of DOT1L and/or SETD2.

Interestingly, we found a significant enrichment of TEAD binding motifs, TEAD4, and YAP1 at regions that gain accessibility in DCI cells (Fig. 4). Such enrichment is unlikely caused by the change in YAP1 phosphorylation and cellular localization since the loss of H3K79me/H3K36me3 has little impact on the levels of total YAP1, phosphorylated YAP1, and the nuclear/cytosolic distribution of YAP1. Since both H3K79me and H3K36me3 are found at enhancers (*31-33, 62, 63*), it is possible that nucleosomes harboring H3K79me and H3K36me3 impede transcription factors such as TEADs from binding to their target enhancers. In addition, DNA methylation may play a role in regulating the recruitment of TEADs to their binding sites. H3K36me3 is known to recruit DNMT3B which methylates DNA on the bodies of active genes (*24, 25*). In addition, DNMT3A has been shown to “read” H3K36me3(*64*). Therefore, determining if H3K36me3 and H3K79me have a synergistic role in regulating DNA methylation levels on gene bodies and enhancers and identifying the role of DNA methylation on the recruitment of YAP-TEAD would be interesting future studies. It is also worth noting that chromatin decompaction and increased recruitment of YAP-TEAD to enhancers in DCI cells could be consequences of hyperactive transcription. Chromatin decompaction and YAP-TEAD recruitment at these enhancers could then feedback to increase transcription levels of target genes. For instance, YAP1 has been shown to interact with NCOA6 (*65*), a subunit of the major enhancer co-activator MLL3/4/COMPASS, and to recruit MLL3/4 and activate YAP1 target genes. The recruitment of MLL3/4/COMPASS to enhancers could decompact chromatin and may further facilitate YAP-TEAD to be recruited to these enhancers. Dissecting the relationship between transcriptional co-activators and YAP-TEAD recruitment at enhancers in the future is also important to understand the mechanisms underlying the synergistic transcription regulatory role of H3K79me and H3K36me3. Interestingly, we did not observe a global increase of chromatin accessibility upon H3K79me/H3K36me3 loss, suggesting that additional factors are involved in regulating chromatin compaction/decompaction at enhancers to safeguard them from hyperactivation. Indeed, transcriptional co-repressor complexes such as LSD1-CoREST and MBD-NuRD are found at many active enhancers (*66-68*). It is possible that H3K79me and H3K36me3 function together with co-repressors as a multi-layered system to safeguard proper enhancer activity. Understanding the relationship between the H3K79me/H3K36me3 synergism and the co-repressor/co-activator network would be very interesting future studies.

Disrupting the binding between YAP and TEADs by verteporfin restores the expression of many misregulated genes in DCI cells (Fig. 5). YAP-TEAD also contributes to chromatin decompaction in DCI cells. These data suggest that the YAP-TEAD axis is partially responsible for the transcription activation caused by the loss of H3K79me/H3K36me3. Our results further suggest a novel approach to target diseases driven by the loss of H3K79me and/or H3K36me3 with YAP-TEAD inhibition. Since DOT1L and SETD2 are both implicated in neurodevelopmental diseases (*12, 69-71*), it would be important to evaluate the impact of YAP perturbation on animal models and patient samples with DOT1L or SETD2 mutations in future studies to determine if targeting YAP-TEAD could potentially benefit neural defects caused by aberrant H3K79me and/or H3K36me3.

In summary, our study unveils a novel synergistic role of H3K79me and H3K36me3 in dampening transcription hyperactivation and safeguarding neural differentiation. Importantly, we identify YAP-TEAD recruitment as a molecular mechanism underlying the transcription misregulation caused by the loss of H3K79me and H3K36me3, providing insight into elucidating the fundamental role of these two conserved histone modifications in gene regulation and understanding human diseases related to the misregulation of them.

## Materials and Methods

### Antibodies

Primary antibodies used in this study were anti-SETD2 (Cell Signaling Technology 80290), anti-H3 (Abcam ab1791), anti-H3K79me1 (Abcam ab2886), H3K79me2 (Abcam ab3594), H3K79me3 (Cell Signaling Technology 74073), H3K36me2 (Cell Signaling Technology 2901), H3K36me3 (Cell Signaling Technology 4909), anti-Tubulin (Developmental Studies Hybridoma Bank E7), anti-HSP90 (Santa Cruz sc-13119), anti-RBBP5 (Bethyl A300-109A), anti-GFAP (Abcam ab7260); anti-TUJ1 (Millipore MAB1637); anti-RNA Pol II (Cell Signaling Technology 14958), anti-RNA Pol II phospho-CTD Ser-5 (Millipore 04-1572), anti-YAP1 (Novus Biologicals NB110-58358), anti-phosphorylated YAP (Cell Signaling Technology 4911), and anti-TEAD4 (Santa Cruz sc-101184). The secondary antibodies used were donkey anti-rabbit IgG HRP (Sigma NA934V), sheep anti-mouse IgG HRP (Sigma NA931V), goat anti-mouse IgG-Alexa Fluor 488 (Life Technologies A11029), and goat anti-rabbit IgG-Alexa Fluor 594 (Life Technologies A32740).

### ESC culture and transfection

V6.5 mouse ESCs were cultured in N2B27-based serum-free medium containing MEK inhibitor PD0325901, GSK inhibitor CHIR99021, and leukemia inhibitory factor (LIF) (Sigma ESG1107) as previously described (*66*). ESCs were transfected with plasmids containing desired guide RNAs (gRNAs) using a Nucleofector (Lonza) as previously described (*72*). ESC clones were picked 10 days after transfection. PCR and Sanger sequencing were used for genotyping the clones. Two independent clones of each genotype have been used in all assays described in this study. The sequences of gRNAs used for generating the SETD2 CI allele were GAGGAGTAGCAGTGCAGTGA (left) and GGTAATAAAGGTATAATCTG (right).

### Histone extraction and analysis

Histones were purified from ESCs based on the procedure previously described (*73*). Briefly, cells were resuspended in ice-cold sucrose buffer and homogenized with a Dounce homogenizer on ice. Pelleted nuclei were then resuspended in ice-cold high-salt buffer and homogenized. Pelleted chromatin was resuspended in 0.2N sulfuric acid, broken apart with a pellet pestle, incubated overnight at 4 °C, and pelleted by centrifugation. Extracted histones were then ethanol precipitated from the supernatant, washed with 70% ethanol, and resuspended in PBS before downstream analysis.

### Neural differentiation

Neural differentiation was performed as previously published (*4*). In brief, embryoid bodies (EBs) were generated from ESCs using the hanging-drop method (*74*). Day 6 EBs were reseeded on 10- cm low-attachment dishes (Sigma) and treated with 1 µM all-trans retinoic acid (Sigma) for 2 days. Day 8 EBs were reseeded on plates coated with poly-L-ornithine (5 µg/mL) and laminin (5 µg/mL), respectively. Upon day 8, EBs were cultured with NeuroCult NSC proliferation medium (StemCell Technologies) supplemented with 10 ng/mL FGF2 (Peprotech) for 21 days. Media was changed every 2 days.

### ChIP with reference exogenous genome (ChIP-Rx) and data analysis

ChIP-Rx was performed as previously described (*66*) with at least two biological replicates from two independent cell clones. In brief, chromatin from fixed ESCs were sheared with an E220-focused ultrasonicator (Covaris) and mixed with 20% of chromatin from HEK293T cells for spike-in normalization. Immunoprecipitation was performed at 4 °C followed by reverse-crosslinking and DNA purification. KAPA HyperPrep Kit (Roche) was used to prepare ChIP-Rx libraries. Libraries were pooled and sequenced on the NovaSeq platform (Illumina) with a read length configuration of 150 bp on each end.

ChIP-Rx mapping and peak calling were performed using the ENCODE ChIP-seq analysis pipeline (https://github.com/ENCODE-DCC/chip-seq-pipeline2). Raw reads were processed with Trim Galore v0.6.6 (https://www.bioinformatics.babraham.ac.uk/projects/trim_galore/) to remove adaptors and low-quality reads with the parameter “-q 25” and then aligned to the mouse mm9 and human hg19 genome assemblies using Bowtie v2.4.4 or BWA v0.7.17 with default parameters (*75, 76*). All unmapped reads, low mapping quality reads (MAPQ < 30), and PCR duplicates were removed using SAMtools v1.14 (*77*) and Picard v2.21.4 (https://broadinstitute.github.io/picard/). The number of spike-in hg19 reads was counted with SAMtools v1.14 and normalization factor alpha = 1e6/hg19_count was calculated. Normalized bigwig was generated with bamCoverage function from deepTools v3.5.1 using scale factors calculated above and reads mapped to the ENCODE blacklist regions were removed using BEDTools v2.17.0 (*78-80*). H3K79me1/2/3 and H3K36me3 peaks were called using Homer and peak annotation was performed with R package ChIPseeker v1.38.0 (*81, 82*). Overlapping and unique peaks were generated using findOverlaps function from R package GenomicRanges v1.46.0 (*83*). For occupancy boxplot representation from the heatmaps, the matrix generated from deepTools v3.5.1 (*80*) was used to calculate the average coverage under each genome region. Pausing indexes were calculated based on the ratio of the RNAPII level (CPM) at promoters defined as -200 bp upstream to 400 bp downstream of TSS and that at the remainder of the entire gene body. ECDF analysis was performed using the ggplot function in R version 4.3.1 and p-values were calculated with a two-sided Kolmogorov-Smirnov test.

### RNA-Seq and data analysis

All RNA-seq experiments were performed with at least two biological replicates from two independent cell clones. RNA was extracted and purified with Trizol reagent (Life Technologies) according to manufacturer’s instructions. RNA was further treated with DnaseI (Sigma) and then purified with RNeasy mini kit (Qiagen). NEBNext rRNA Depletion Kit (New England BioLabs) and NEBNext Ultra II Directional RNA Kit (New England BioLabs) were used to deplete ribosomal RNA and prepare RNA-seq libraries, respectively. Libraries were pooled and sequenced on the NovaSeq platform (Illumina) with a read length configuration of 150 bp on each end.

Raw reads from RNA-seq were trimmed as described in ChIP-Rx and then aligned to the mm9 genome assembly using STAR v2.5.1 with default parameter (*84*). Reads were normalized to total read counts per million (cpm) and visualized as bigwig-formatted coverage tracks using deepTools v3.5.1 (*80*). Gene expression quantification was performed with HTseq v0.11.1 (*85*). Differential expression analysis was performed using R package EdgeR v3.40.2 (*86*). Significant differentially expressed genes were filtered out with Benjamini-Hochberg-adjusted *p* values less than 0.01 and log fold change larger than |1|. Customed R scripts were used to generate correlation plots. Boxplots were generated by ggboxplot function from package ggpubr v0.6.0 (https://rpkgs.datanovia.com/ggpubr/). GSEA analysis was performed by R package clusterProfiler v4.10.1 (*87*).

### ATAC-seq and data analysis

ATAC-seq was performed as described previously (*88*). In brief, cells were washed with PBS and lysed with ATAC lysis buffer. 50K nuclei were tagmented at 37 °C for 30 minutes. Tagmented DNA was then cleaned up with the MinElute Reaction Clean-up Kit (Qiagen) and subject to library preparation. Libraries were pooled and sequenced on the NovaSeq platform (Illumina) with a read length configuration of 150 bp on each end.

Raw reads from ATAC-seq were trimmed as described in ChIP-Rx and aligned to the mm9 genome assembly using Bowtie v2.4.4 (*75*) with parameter “-N 1 -L 25 -X 2000 --no-mixed --no-discordant”. Mitochondrial reads and PCR duplicates were then filtered using SAMtools v1.12 (*77*) and Picard Tools v2.25.5 (https://broadinstitute.github.io/picard/). Finally, the reads were shifted to compensate for the offset in the tagmentation site relative to the Tn5 binding site using the alignmentSieve function of deepTools v3.5.1(*80*) with the ‘--ATACshift’ option. Reads were normalized to total read counts per million (cpm) and visualized as bigwig-formatted coverage tracks using deepTools v3.5.1 (*80*). ATAC-seq peaks were called using MACS2 v2.2.7.1 (*89*), peaks called from replicates in WT and mutant cells were merged by bedTools v2.30.0 (*79*). A raw-count matrix of merged peaks was generated using featureCounts v2.0.2 (*90*) and then differential peak analysis was performed using DEseq2 v1.32.0 (*91*). Motif analysis was performed using findMotifsGenome.pl function of HOMER (*81*).

## Acknowledgements

We thank the Cao lab members for their helpful suggestions and discussions. We thank Yalu Zhou for bioinformatic analysis support. This study was supported in part by grants R00HD094906, P30CA043703, and R35GM150668 from the National Institutes of Health to K.C.

## Author Contributions

K.C. conceived the study and designed the research. E.W.C., C.Z., S.M.N., Y.J., A.H., and K.C. performed experiments. E.W.C., C.Z., S.M.N., Y.J., A.H., J.C., P.G., and K.C. analyzed the data. FX.C. and F.J. provided resources for bioinformatic analysis. K.C. and E.W.C. wrote the manuscript.

## Figure Legends

**Figure S1.**
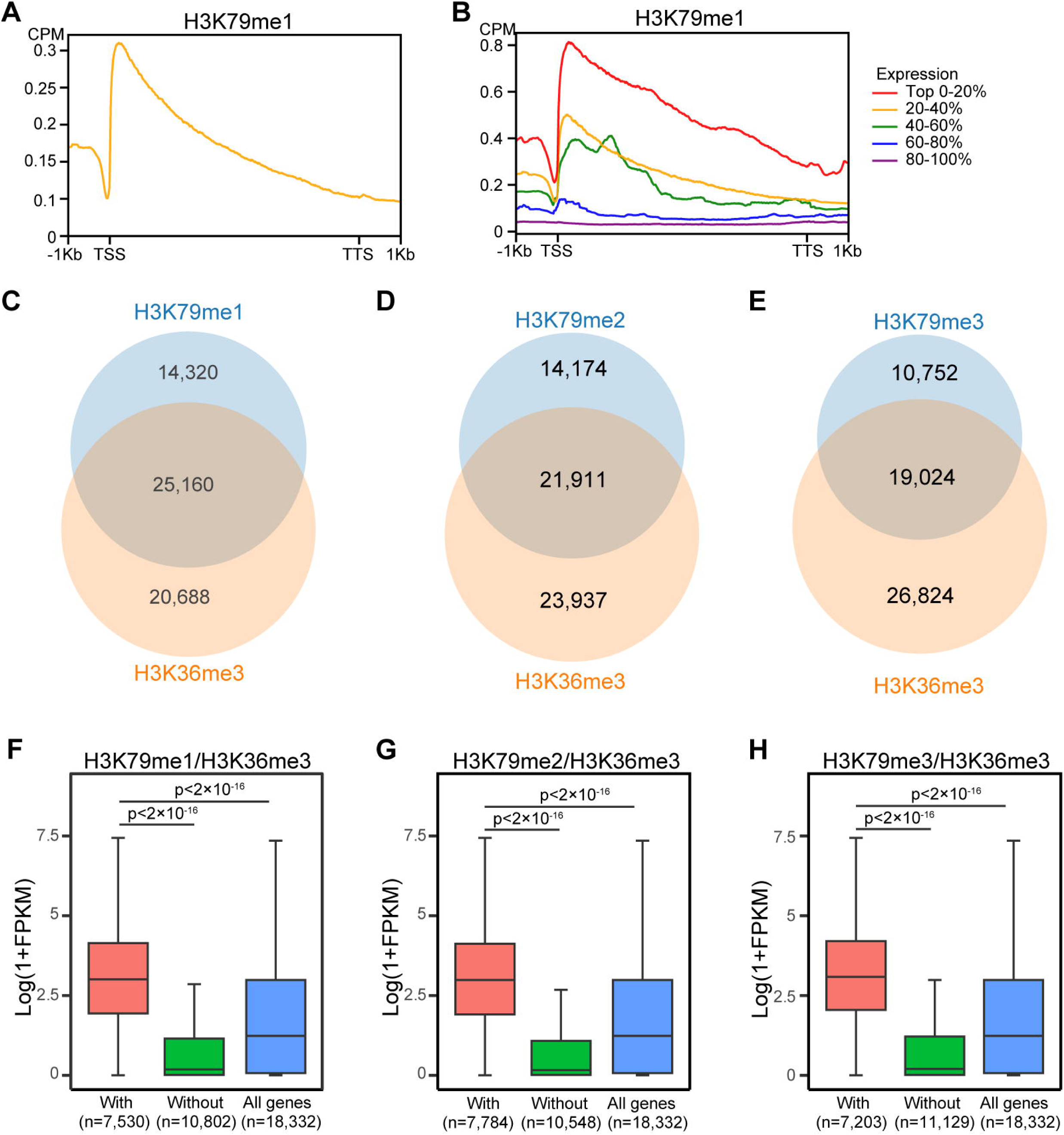
Genome-wide colocalization of H3K9me and H3K36me3. (A) Metaplots of average ChIP-Rx signals of H3K79me1 on all protein-coding genes (n=18,332) in ESCs. (B) Metaplots showing the signals of H3K79me1 on all protein-coding genes categorized into five groups based on their expression levels. (C-E) Venn diagrams showing the colocalization of H3K79me1 (C), H3K79me2 (D), and H3K79me3 (E) peaks with H3K36me3 peaks, respectively. (F-H) Box plot analysis of the expression levels of genes with and without H3K79me1/H3K36me3 (F), H3K79me2/H3K36me3 (G), and H3K79me3/H3K36me3 (H) peaks. The number of genes in each category is labeled.

**Figure S2.**
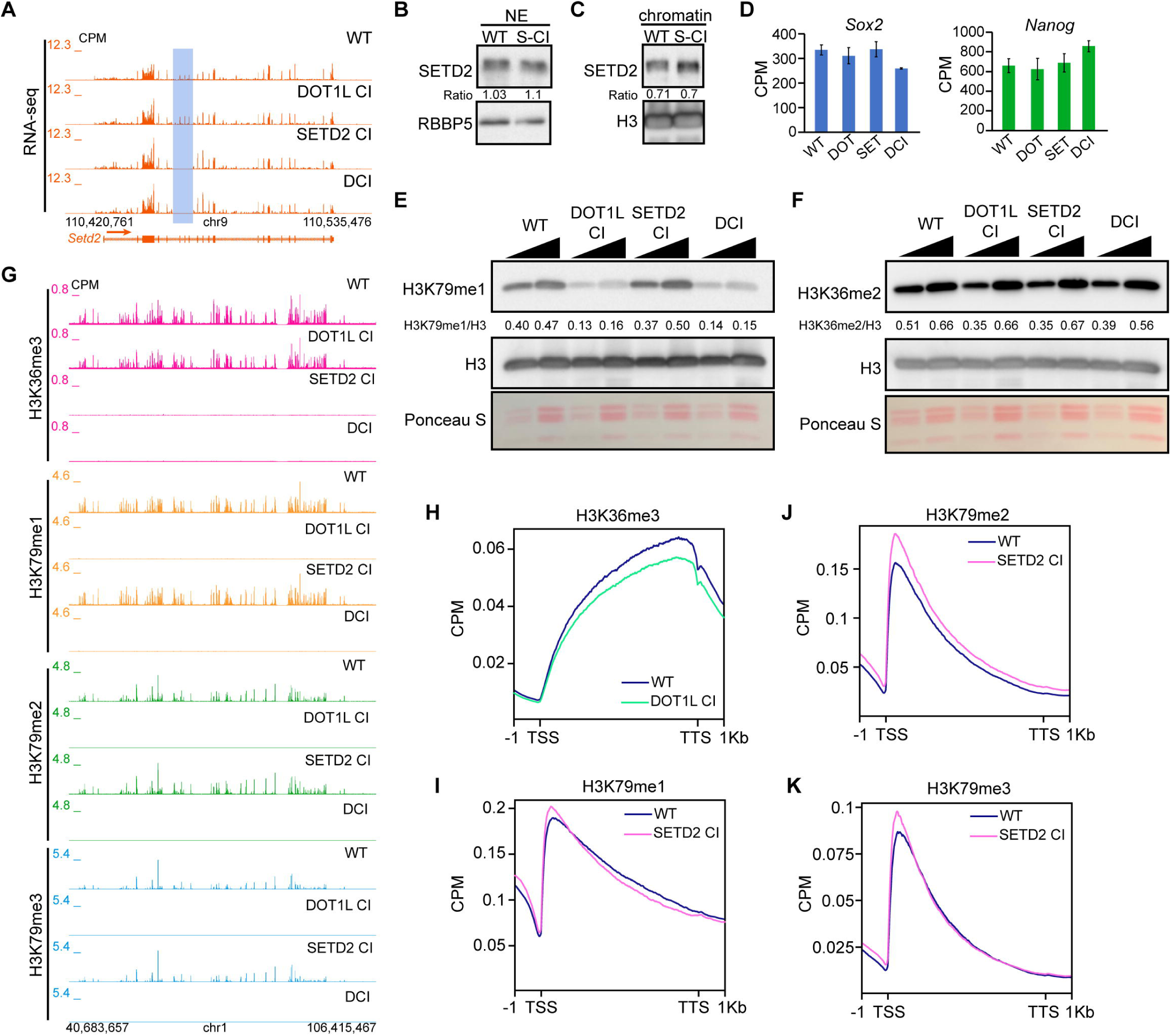
Generation and characterization of SETD2 CI and DCI cells. (A) The genome browser view of RNA-seq signals at the *Setd2* gene in WT, DOT1L CI, SETD2 CI, and DCI ESCs. Deleted regions in SETD2 CI and DCI cells are highlighted in blue. (B-C) Western blotting of SETD2 in nuclear extract (NE) (B) and chromatin (C) fractions of WT and SETD2 CI (S-CI) cells. Relative SETD2 levels were quantified by normalizing with the corresponding controls and labeled below the SETD2 blot. (D) RNA-seq signals of *Sox2* (left) and *Nanog* (right) genes in WT, DOT1L CI (DOT), SETD2 CI (SET), and DCI cells. (E-F) Western blotting analysis indicating the level of H3K79me1 (E) and H3K36me2 (F) in WT, DOT1L CI, SETD2 CI, and DCI ESCs. 1 and 2 μg extracted histones were loaded for each sample. (G) Genome browser view of H3K36me3, H3K79me1, H3K79me2, and H3K79me3 ChIP-Rx signals in WT, DOT1L CI, SETD2 CI, and DCI cells. (H) Metaplot analysis showing the level of H3K36me3 in WT and DOT1L CI cells on all protein-coding genes. (I-K) Metaplots showing the level of H3K79me1 (I), H3K79me2 (J), and H3K79me3 (K) in WT and SETD2 CI cells on all protein-coding genes.

**Figure S3.**
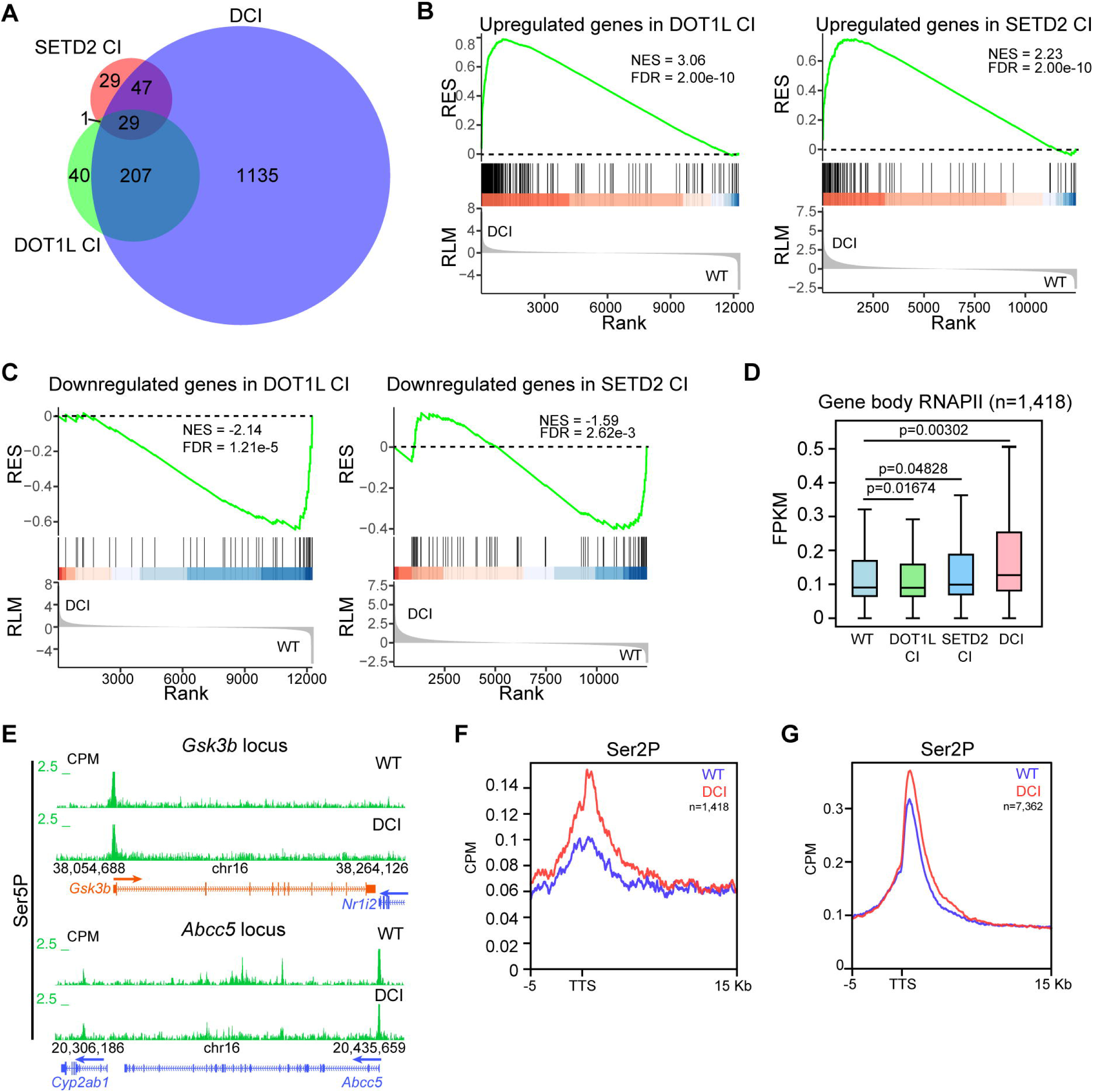
H3K79me and H3K36me3 synergistically regulate gene expression and transcription elongation. (A) Venn diagram showing the overlap of upregulated genes in DOT1L CI, SETD2 CI, and DCI cells compared with WT cells. (B-C) GSEA analysis of upregulated (B) and downregulated (C) genes in DOT1L CI (left) and SETD2 CI (right) comparing DCI and WT cells. RES: running enrichment score; RLM: ranked list metric; NES: normalized enrichment score; FDR: false discovery rate. (D) Box plot analysis of RNAPII levels at gene bodies of the 1,418 DCI upregulated genes (red genes in Fig. 3C) in WT, DOT1L CI, SETD2 CI, and DCI cells. FPKM: spike-in normalized fragments per kilobase per million mapped reads. (E) Genome browser view of Ser5 phosphorylated RNAPII (Ser5P) ChIP-Rx signals at *Gsk3b* (up) and *Abcc5* (down) loci in WT and DCI cells. (F-G) Metaplot analysis of Ser2 phosphorylated RNAPII (Ser2P) ChIP-Rx levels at -5Kb and +15Kb around TTS of 1,418 DCI upregulated genes (F) and 7,362 RNAPII positive genes (G) in WT and DCI ESCs.

**Figure S4.**
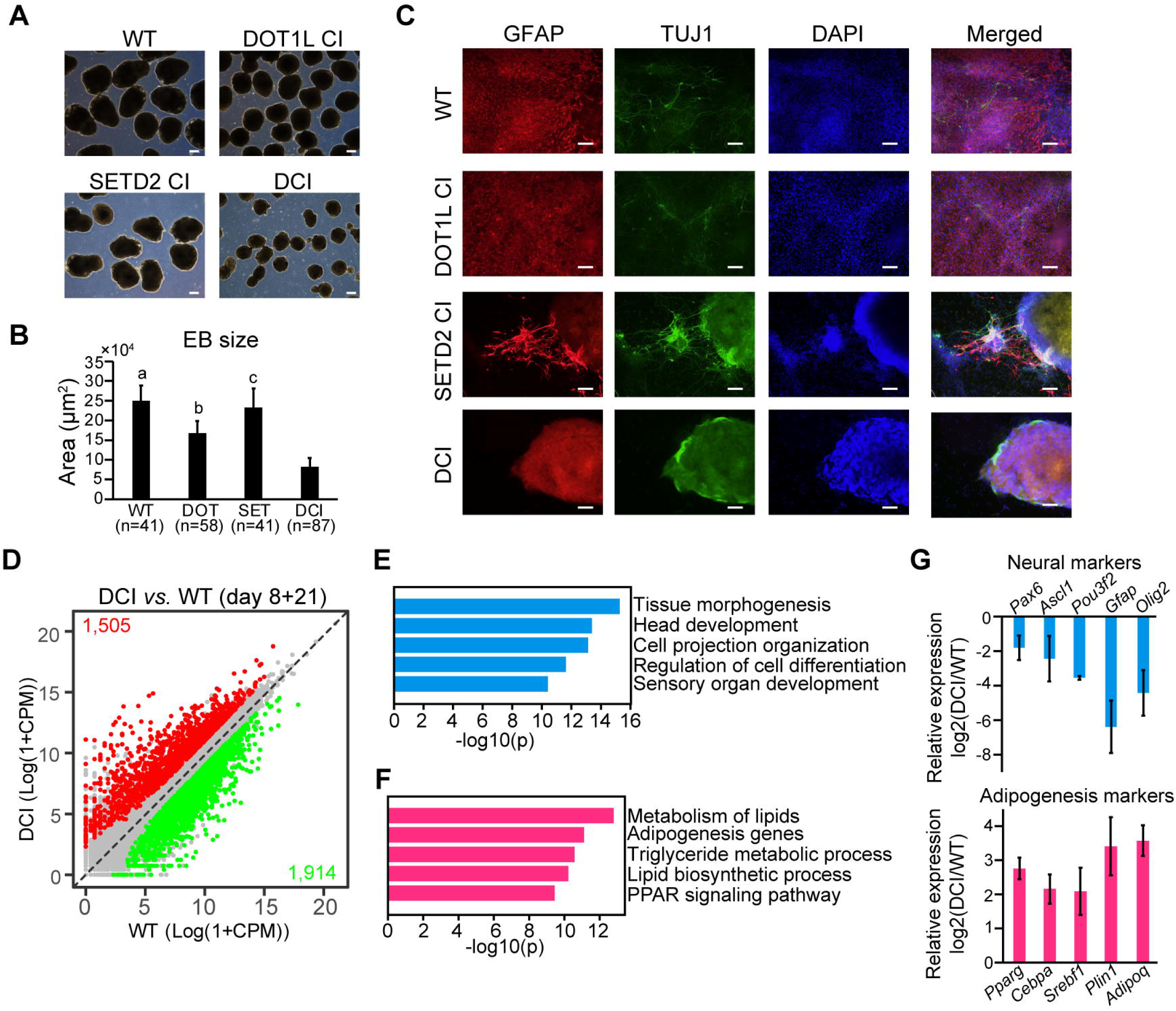
H3K79me and H3K36me3 synergistically regulate neural differentiation. (A) Phase-contrast images of WT, DOT1L, SETD2CI, and DCI EBs at day 6. Scale bar: 250µm. (B) Quantification of EB sizes in (A). Data are presented as mean values ± standard deviation (SD). n = 3 biologically independent experiments. a: p= 4.36×10^-31^; b: p=3.68×10^-32^; c: p= 6.9×10^-23^. P-values were calculated using a two-sided student’s t-test comparing EB sizes of each group with those of DCI EBs. (C) Day 8+21 neural differentiation cultures of WT, DOT1L CI, SETD2 CI, and DCI cells were immunostained with GFAP- and TUJ1-specific antibodies to show respective astrocytes and neurons in culture. Nuclei were stained with DAPI. Scale bar: 100µm. (D) Correlation plots of RNA-seq data between day 8+21 neural differentiation cultures of DCI and WT cells. Statistical significance was determined by a two-sided Wald test and p-values were corrected for multiple testing using the Benjamini-Hochberg method. Significantly upregulated genes (log2 fold change >1, adjusted p<0.01) are highlighted in red while downregulated (log2 fold change <-1, adjusted p<0.01) genes are highlighted in green. The number of up- and downregulated genes are listed on the plots. (E-F) Gene ontology (GO) analysis of top 300 downregulated (E) and upregulated (F) genes in DCI cells after 29 days of neural differentiation. The 5 most significant GO terms are shown in each plot. (G) Fold change of RNA-seq signals (CPM) in DCI over WT neural differentiation culture is shown for neural (top) and adipogenesis (bottom) markers. Data are presented as mean values ± SD.

**Figure S5.**
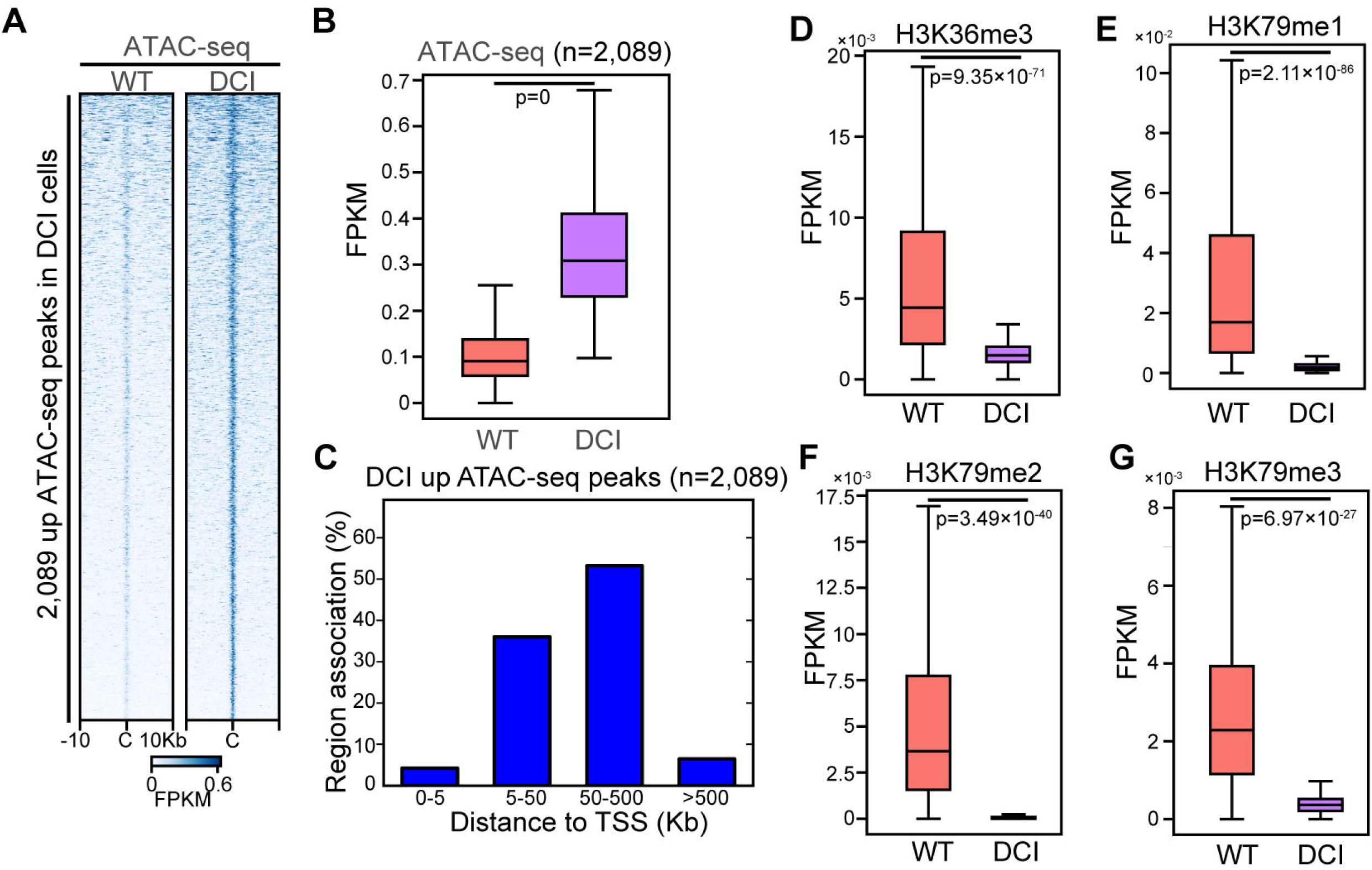
H3K79me/H3K36me3 loss leads to gained chromatin accessibility at enhancers. (A) Heatmaps showing the levels of ATAC-seq signals at DCI-upregulated ATAC-seq peaks in WT and DCI cells. Data are sorted by descending ATAC-seq signals in DCI cells. (B) Box plot analysis indicating the levels of ATAC-seq signals at DCI-upregulated ATAC-seq peaks in WT and DCI cells. (C) Distance of DCI upregulated ATAC-seq peaks to TSS are shown by bar plots. Analysis was performed using GREAT v4.0.4(*92*). (D-G) Box plot analysis indicating the levels of H3K36me3 (D), H3K79me1 (E), H3K79me2 (F), and H3K79me3 (G) at 2,089 DCI upregulated ATAC-seq peaks in WT and DCI cells.

**Figure S6.**
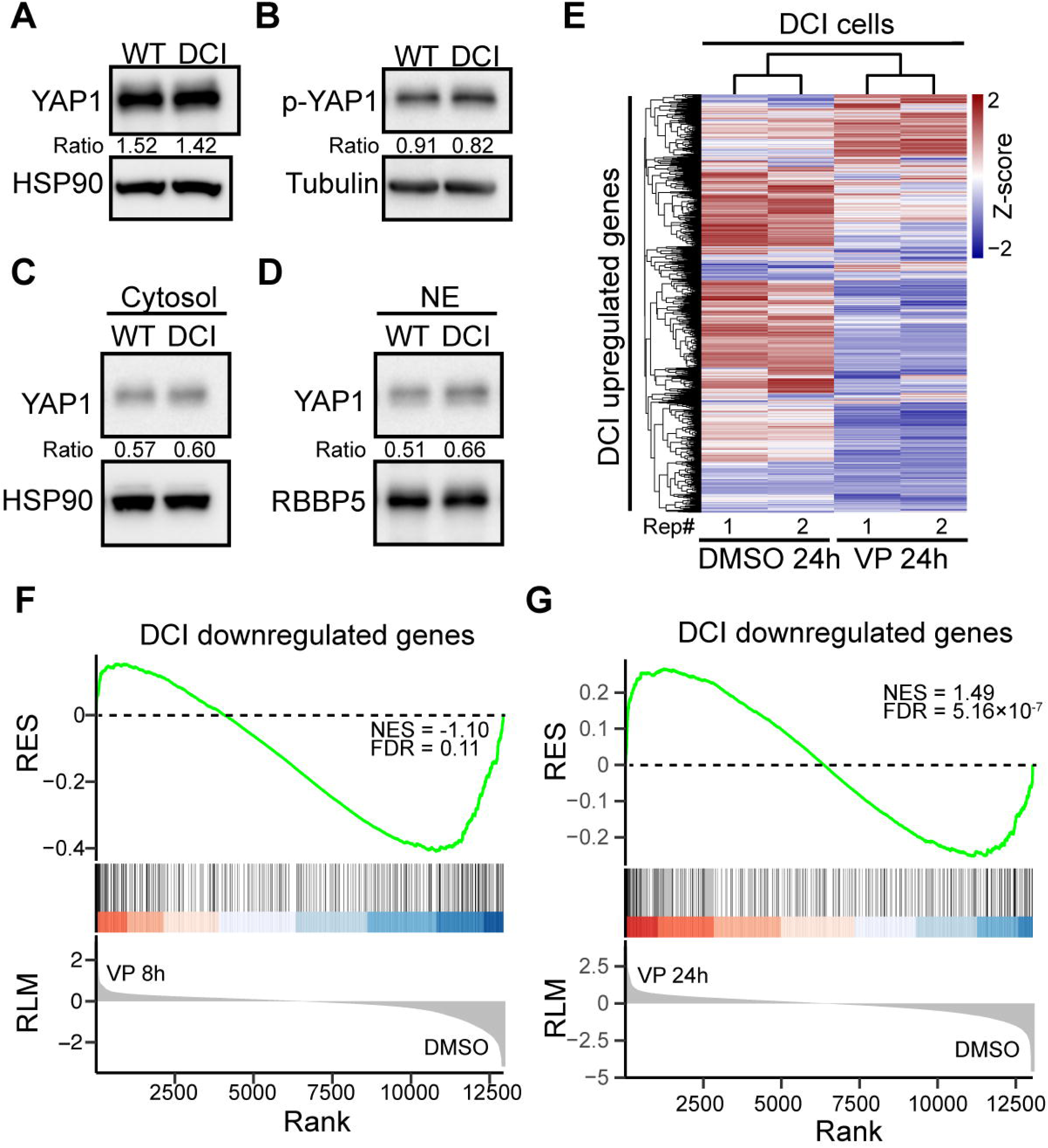
The role of YAP-TEAD in gene misregulation caused by the H3K79me/H3K36me3 loss. (A-B) Western blotting analysis of YAP1 (A) and phosphorylated YAP1 (p-YAP1) (B) using whole cell lysate of WT and DCI cells. (C-D) Western blotting analysis of YAP1 in cytosolic (C) and nuclear (D) fractions of WT and DCI ESCs. NE: nuclear extract. (E) Hierarchical clustering analysis of expression levels of the 1,418 DCI upregulates genes comparing DCI cells treated with DMSO or VP for 24 hours. Z-scores were used to generate the heatmap. Numbers below the heatmap denote the 2 biological replicates of each treatment condition. (F-G) GSEA analysis of genes downregulated in DCI cells (blue genes in Fig. 3c) comparing 8-hour (F) and 24-hour (G) VP- and DMSO- treated DCI cells. RES: running enrichment score; RLM: ranked list metric; NES: normalized enrichment score; FDR: false discovery rate.

